# Binding Dynamics of Disordered Linker Histone H1 with a Nucleosomal Particle

**DOI:** 10.1101/2020.11.25.398180

**Authors:** Hao Wu, Yamini Dalal, Garegin A. Papoian

## Abstract

Linker histone H1 is an essential regulatory protein for many critical biological processes, such as eukaryotic chromatin packaging and gene expression. Mis-regulation of H1s is commonly observed in tumor cells, where the balance between different H1 subtypes has been shown to alter the cancer phenotype. Consisting of a rigid globular domain and two highly charged terminal domains, H1 can bind to multiple sites on a nucleosomal particle to alter chromatin hierarchical condensation levels. In particular, the disordered H1 amino- and carboxyl-terminal domains (NTD/CTD) are believed to enhance this binding affinity, but their detailed dynamics and functions remain unclear. In this work, we used a coarse-grained computational model AWSEM-DNA to simulate the H1.0b-nucleosome complex, namely chromatosome. Our results demonstrate that H1 disordered domains restrict the dynamics of both globular H1 and linker DNA arms, resulting in a more compact and rigid chromatosome particle. Furthermore, we identified regions of H1 disordered domains that are tightly tethered to DNA near the entry-exit site. Overall, our study elucidates at near atomic resolution the way the disordered linker histone H1 modulates nucleosome’s structural preferences and conformational dynamics.

## Introduction

Eukaryotic chromatin consists of histone proteins and DNA [1]. Four types of histone proteins, namely H2A, H2B, H3, and H4, are assembled as an octamer (two H2A-H2B dimers and one H3-H4 tetramer), and wrapped around by ~147 base pairs (bp) of DNA [2]. These elementary repeating units, called nucleosomes, are connected by short linker DNA and the linker histone protein H1, forming a “zigzagging ladder” architecture. These nucleosomal arrays are folded in several hierarchical levels to form chromatin in the nucleus [3].

The fundamental mechanisms of this intricate folding process, still, remain poorly understood. How does chromatin change its condensation levels rapidly and accurately, so that a gene can be precisely transcribed while being well protected from DNA damage [4]? The linker histone H1 is believed to play an important role in the fast dynamics of chromatin folding [5–8]. As a developmentally regulated protein, H1 has a family of variants that are specific to distinct species or tissues [9]. All these H1 variants consist of three parts: a short (20 - 40 amino acids (AA)) disordered amino-terminal domain (NTD), a highly-conserved globular domain (~80 AA) with rigid structure, and a long (100-125 AA) highly disorganized carboxyl-terminal domain (CTD) with varying lysine and arginine contents, making various binding affinities possible amongst the variants [10–12]. The high mobility of H1 when bound to a nucleosome is directly related to many biological processes [13–17]. Meanwhile, H1 depletion levels are found to alter global nucleosome spacing and local chromatin compaction [18,19]. H1 variants are modified and regulated by various post-translational modifications (PTMs), which in turn are thought to confer distinct chromatin structures [20].

The first crystal structure of the linker histone globular domain at high-resolution (2.6 Å) discovered its “winged-helix” folding motif over two decades ago [21]. However, the high-resolution structural insights of the linker histone-nucleosome complex, referred as to chromatosome, remained elusive until five years ago. Consequently, compared with the four core histone proteins, the structure and dynamics of linker histones are less studied and understood. Interestingly, various H1 subtypes are found to bind on different locations of the nucleosome under different experimental conditions [22]. *Drosophila* H1 [23] and human H1.4 [24] bind to nucleosome near dyad – the center bp defining pseudo-two-fold symmetry axis of the nucleosome – in an asymmetric manner, while chicken linker histone H5 [25,26], *Xenopus Laevis* H1.0b, and human H1.5 [27] bind right on the center of the dyad symmetrically. Zhou *et al*. proposed these “on-dyad” and “off-dyad” binding modes will lead to distinct chromatin condensation levels [25]. A series of atomistic computational studies revealed the conformational selection mechanism of these H1-nucleosome binding modes, as well as the effects of sequence and PTMs [28–30]. Further cryo-EM studies [24,31] and large-scale simulations with ~1 nm spatial resolution [32,33] (i.e. mesoscale simulations) of nucleosomal arrays explored how the linker histone’s subtype and concentration determine inter-nucleosome relative positions and resulting distinct chromatin geometries. More experimental studies also discovered other important biological functions of H1 variants besides altering chromatin structure, such as regulating gene expression [18], generating epigenetic heterogeneity within tumor cells [34], and directly inhibiting transcriptions [35].

Most of the previous studies on linker histones have focused on the globular domain, while the structure and dynamics of the N- and C-termini remain poorly understood, in part because of their intrinsically disordered nature. Recent studies reveal that H1 NTD and CTD are extremely disordered, even when bound with other proteins [36] or DNA [37]. Their disordered nature, in turn, imparts unique functions of liquid-like glue to promote chromatin folding [38]. In particular, the long and highly basic H1 CTD was found to be more essential for the linker histone to bind onto nucleosome with high affinity compared to the NTD [8,39]. Similar to the core histone tails, which have been extensively studied by computer simulations [40–42], H1 CTD is also highly disordered and bound to DNA, but much longer than the ~15-40 AA-long core histone tails. A recent cryo-EM experimental study validated the function of CTD in stabilizing the H1-nucleosome complex by primarily binding to one of the linker DNA arms [27]. A recent comprehensive study using cryo-EM, NMR, and MD simulation of three human H1 isoforms further confirmed H1 CTD’s stabilizing effect on DNA dynamics and discovered new binding dynamics between H1 and core histone tails [43]. FRET experiments [44,45] and mesoscale simulations [32] demonstrated a variety of H1 CTD conformations depending on the environment of nucleosome arrays and linker DNA length. Furthermore, a series of recent computational studies used both atomistic molecular dynamics simulations and mesoscale Monte Carlo simulations to investigate the H1 disordered domains’ conformation and dynamics in one and multiple chromatosome particles, as well as the regulatory roles of H1 phosphorylation and disorder-to-order transition on the nucleosome asymmetry and chromatin large-scale organization [46,47].

In this work, we studied the structure and dynamics of an H1 variant, *Xenopus Laevis* H1.0b, in complex with a nucleosome particle, via molecular dynamics simulations (we use “H1” below to refer to the *Xenopus Laevis* H1.0b in all the subsequent parts of this study, unless otherwise specified). We used a coarse-grained protein model called AWSEM [48], in combination with recently developed AWSEM-IDP [49] for representing disordered domains, and a coarse-grained DNA force field 3SPN.2 [50], to simulate this large protein-DNA complex, the chromatosome. To analyze separate roles of H1 globular and disordered domains in determining chromatosome structure and dynamics, we simulated three combinations of H1-nucleosome systems: (1) nucleosome alone with linker DNA arms; (2) nucleosome with linker DNA arms and the H1 globular domain (GH1); (3) nucleosome with linker DNA arms and the full-length H1 (see **Figure 1**). Our computational studies showed that the H1 disordered domains restrict conformationally GH1 and linker DNA, shedding new light on H1’s role in chromatosome compaction and chromatin condensation.

**Figure 1:**
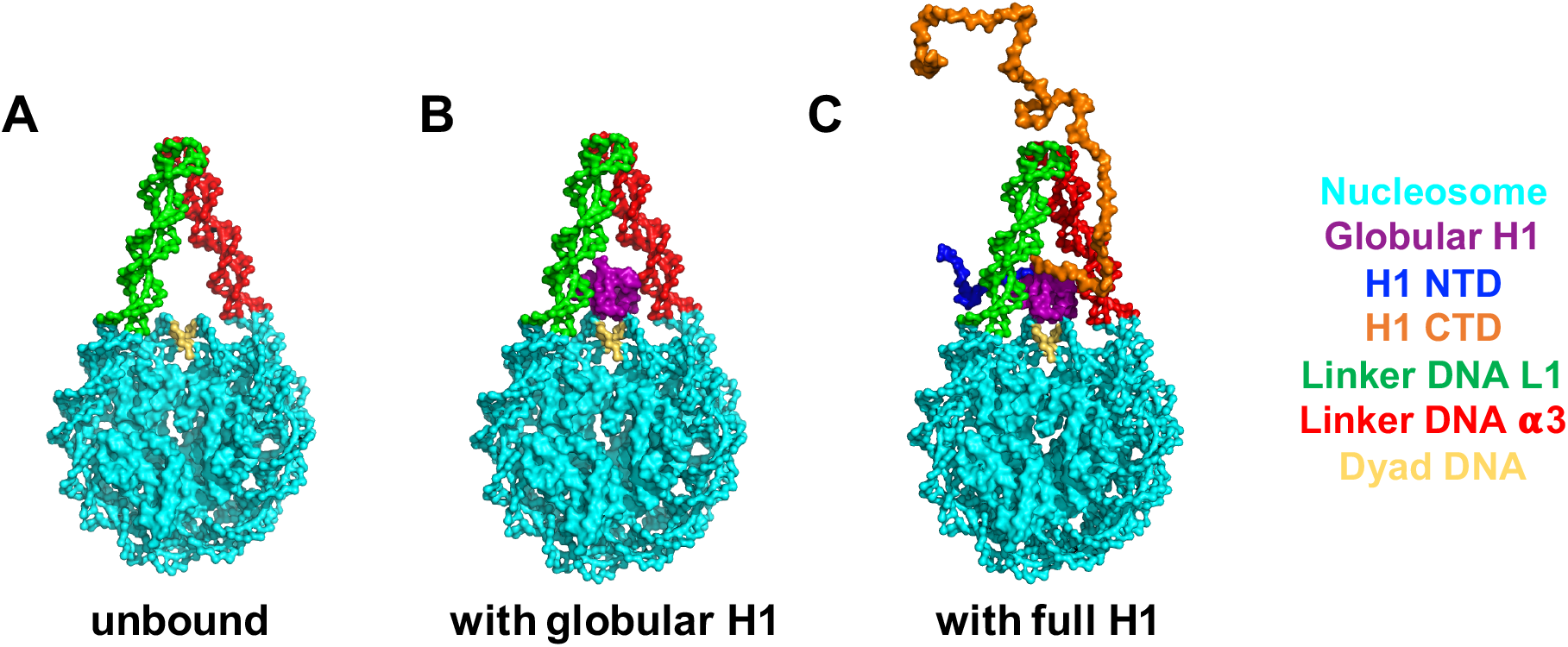
The molecular systems simulated in this study. From left to right: unbound nucleosome without H1 (A), globular H1-nucleosome (B) and full-length H1-nucleosome (C). All the models are based on a recent X-ray crystal structure (PDB: 5NL0) of the *Xenopus Laevis* H1.0b-nucleosome.

## Results

### H1 disordered domains confine GH1 dynamics

To quantify how the presence of H1 disordered domains affects the dynamics of GH1, we tracked the trajectories of GH1’s COM in our simulations (**Figure 2**). The trajectories in the absence of H1 disordered domains demonstrate that GH1 is very dynamic (**Figure 2A**). Starting from the initial position on dyad, GH1 not only swings near the dyad but also may drift away from the nucleosome. By contrast, the presence of H1 NTD/CTD significantly restricts GH1’s range of activity (**Figure 2B**). As a part of the highly basic and disordered full-length H1, the globular domain still explores multiple conformations but all adjacent to the nucleosome. We also computed the radius of gyration (*R*_*g*_) of GH1 COM trajectories to measure their average range of motion. The *R*_*g*_ without H1 NTD/CTD (23.5 Å) is larger than that with H1 NTD/CTD (20.9 Å), further validating H1 disordered domains’ inhibitory role on GH1 dynamics.

**Figure 2:**
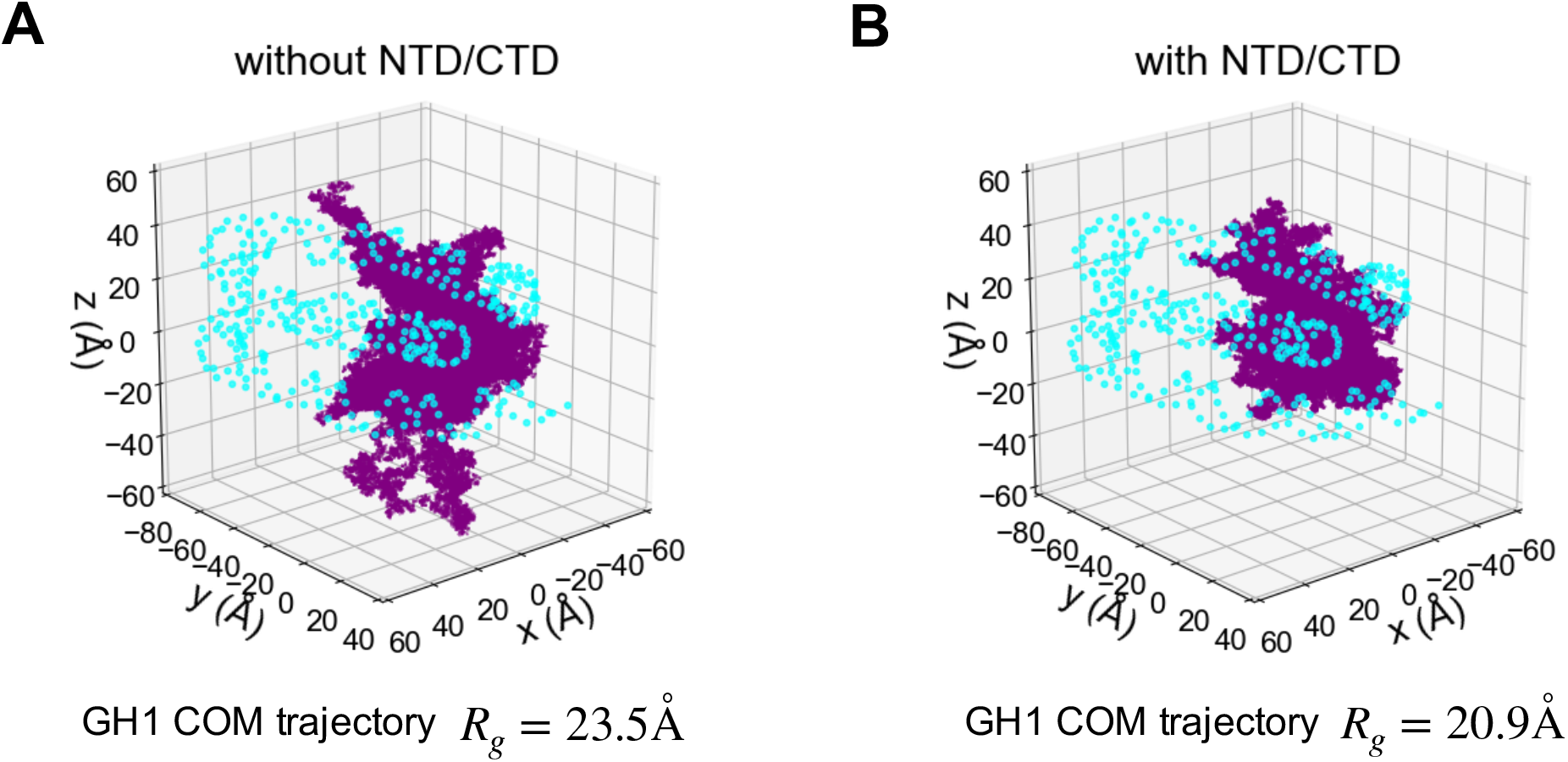
GH1 dynamics is constrained by H1 NTD/CTD. The trajectories of GH1 COM in the H1-nucleosome system without H1 NTD/CTD **(A)** and with H1 NTD/CTD **(B)** are plotted as purple dots. The DNA beads are represented as cyan circles. The core histone octamer and H1 NTD/CTD are not shown for clarity.

The scatter plots of trajectories serve as an overview of GH1 conformations and dynamics, but the preferred GH1-nucleosome binding locations are not clearly identifiable. Therefore, we set up a three-dimensional reference coordinate system (*x, y, z*), and define the location of GH1 COM relative to the nucleosome core particle’s COM in spherical coordinates (*r*, *θ*, *ϕ*) (see **Figure 3A** for detailed definition). Then we computed 2D histograms of all three spherical coordinates to identify the most probable GH1 binding modes. Here we found the effects brought by H1 disordered domains are best represented by the (*ϕ*, *r*) histogram (**Figure 3B-C**, the other two histograms of (*θ*, *ϕ*) and (*θ*, *r*) are shown in Figure S5). For GH1 without the disordered domains, the (*ϕ*, *r*) distribution, especially *ϕ* values, is very dispersed (**Figure 3B**). This means GH1 can almost freely rotate around the vertical *y*-axis across the dyad, whereas in the full-length H1-nucleosome simulations, we found the 2D distribution is more concentrated (**Figure 3C**). Almost all the GH1 conformations with H1 disordered domains have positive *ϕ* values, indicating that GH1 is only allowed to be located on one side of the nucleosomal disk. This analysis further demonstrates that the disordered domains reduce the variability of GH1’s binding modes.

**Figure 3:**
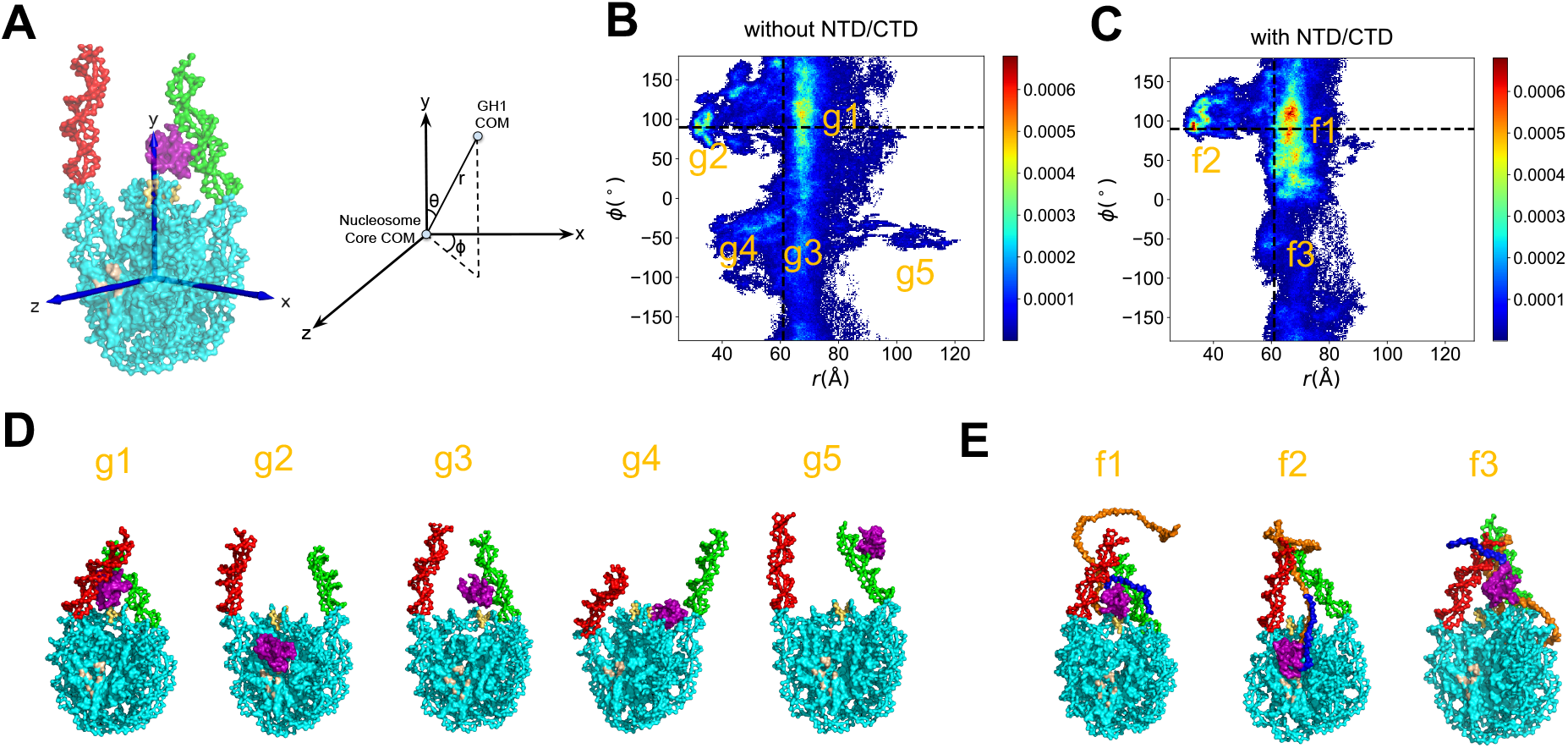
GH1-nucleosome relative conformations. **(A)** Definition of the 3D reference coordinate system and GH1 COM spherical coordinates (*r*, *θ*, *ϕ*). The 2D histogram of (*ϕ*, *r*) for GH1-nucleosome **(B)** and full-length H1-nucleosome **(C)** are shown as heat maps. The color bar from blue to red indicates a probability from low to high. The *ϕ* and *r* values at the beginning of simulations are labeled as black dashed lines. The representative snapshots of all the major basins labeled in **(B-C)** are shown in **(D)** and **(E)**. The color code for different parts of molecules is the same as **Figure 1**, except for the gray region for the histone acidic patch.

### H1 disordered domains restrict GH1-nucleosome interaction sites

Besides describing GH1 dynamics, the 2D histograms in **Figure 3** also identify the preferred GH1-nucleosome binding modes. Here we found five major basins (g1, g2, g3, g4, g5) for GH1-nucleosome and three (f1, f2, f3) for full-length H1-nucleosome. The representative snapshots of each basin are shown in **Figure 3D-E** (see Table S2 and Supporting Information for definitions and population percentages of each basin, and the detailed procedure to select representative snapshots). In two major conformations g1 and g3 of H1-nucleosome without disordered domains, GH1 is still located near the dyad DNA minor groove (labeled as yellow). In basins g2 and g4, however, GH1 tends to slip off the dyad region and move towards the histone octamer (see Supplemental Movie 1 to demonstrate this behavior). With a small probability (~0.9%), GH1 even moves to the tip of one linker DNA arm to escape away from the dyad in basin g5. Meanwhile, four out of these five major basins (25.4%) have very different *ϕ* and *r* values from the on-dyad initial binding mode (black dashed lines). For full-length H1-nucleosome, two out of three basins (f1 and f3) are located more proximal to the initial binding mode. This contrast clearly shows that the presence of H1 NTD/CTD shifts the preferable GH1 binding positions closer to the dyad. Similar to g2, there is also an unignorable basin f2 representing a far-away-from-dyad binding conformation even with NTD/CTD. One possible reason might be the competitive electrostatic attraction from the acidic patch region on histone core (shown as gray in **Figure 3D-E**, also see Figure S6 for snapshots with a zoomed-in view). Overall, this result demonstrates that H1 disordered domains tend to stabilize the GH1’s position near the DNA entry-exit site.

Previous experiments have found that various types of linker histones recognize nucleosomal DNA via different parts of their “winged-helix” folding motif [25,27]. To further probe the DNA-binding preferences of these secondary structure elements, we divided GH1 and DNA into several regions, and computed the contact probability of two arbitrary beads belonging to a certain pair of regions (**Figure 4**). For the GH1-nucleosome without NTD/CTD, the main regions of H1 in contact with dyad DNA are L1/β1 and β2 (**Figure 4B**, 2nd row). By contrast, when NTD/CTD are present, α2 and α3 helical regions form stronger contacts to the dyad DNA (**Figure 4C**, 2nd row). These two helices also have more residues proximal to the dyad DNA in the X-ray crystal structure [27] (labeled with purple stars), suggesting that our results with NTD/CTD have a similar near-dyad binding pattern compared to the experimental results. On the other hand, GH1’s binding probabilities with the α3- and L1-linker DNA (**Figure 4B-C**, 1st, 3rd row) are mostly low with or without the disordered domains. But when NTD/CTD are present (**Figure 4C**, 1st row), the α3 helix forms significantly more contacts with the α3 DNA, consistent with the crystal structure as well. Our contact map analyses provide a detailed description of the GH1-DNA interface in the presence of H1 NTD/CTD: GH1 recognizes nucleosome mainly through dyad DNA by α2/α3 helix. But instead of being located in the exact dyad center, GH1 is tilted towards α3 DNA. The absence of H1 disordered domains, on the contrary, changes this stable interaction network and makes GH1 more flexible.

**Figure 4:**
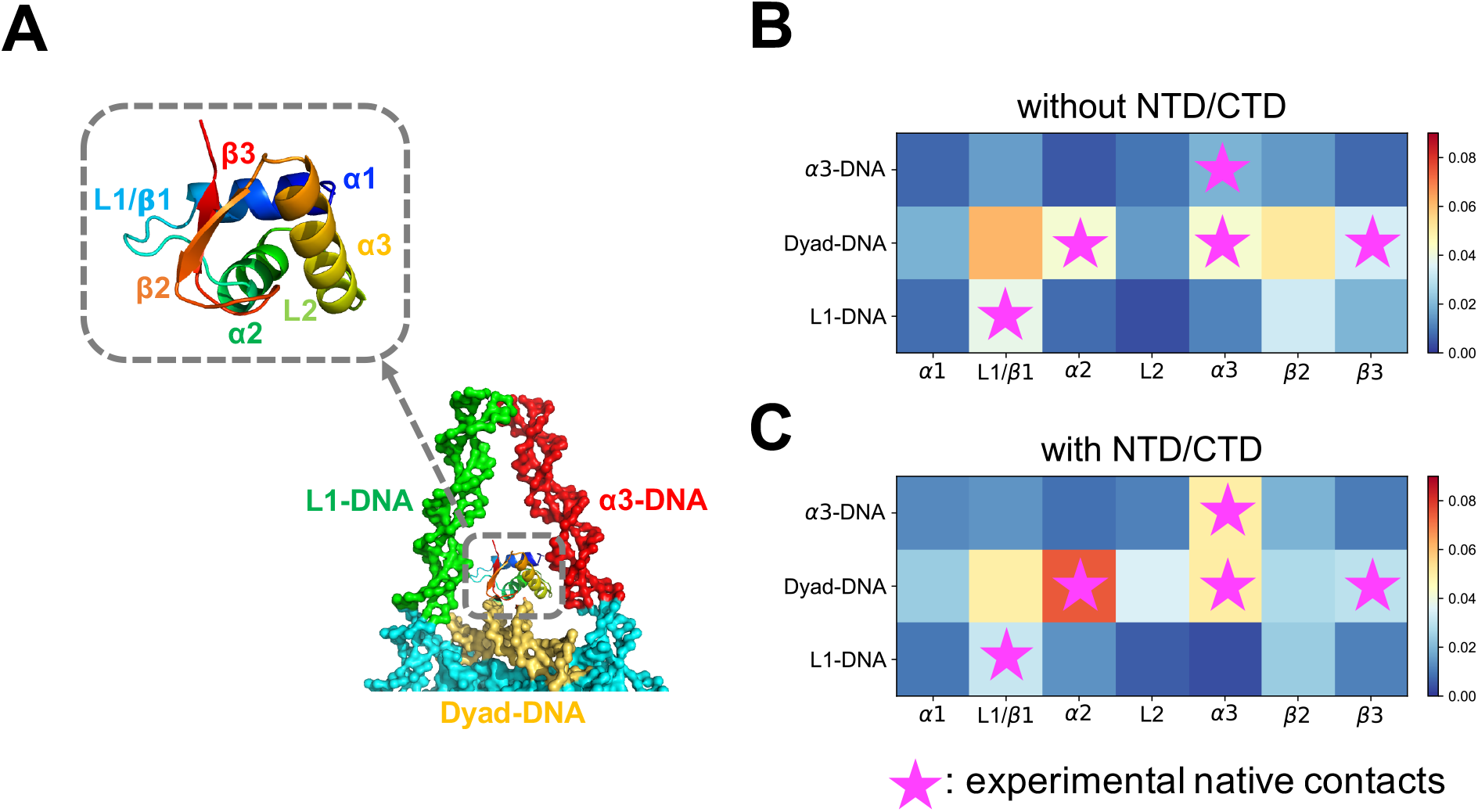
H1 disordered domains regulate and stabilize GH1-DNA binding interface. GH1 and nearby DNA structures are represented in **(A)**, where colors represent different protein or DNA regions. **(B-C)** GH1-DNA contact maps without and with H1 NTD/CTD. Horizontal and vertical axes represent GH1 and DNA regions respectively. Color code from blue to red indicates contact probability of a protein-DNA beads pair in this region from low to high. Purple stars represent the experimental “native contact regions”, where more than two residues in this GH1 region are close to a DNA region.

### H1 globular and disordered domains converge linker DNA

The geometry of linker DNA arms is another important feature to evaluate chromatosome structural compaction. Previous cryo-EM experiments [27] show that H1 induces a more compact and rigid nucleosome conformation by keeping the linker DNA arms convergent or “closed”. Here we computed angles α and β to quantify the linker DNA’s geometry parallel and perpendicular to the nucleosomal disk. We found the α angle distribution of full-length H1-nucleosome (29.3° ± 14.8°) shifts toward higher values compared to the unbound results (15.6° ± 17.7°) (**Figure 5A**), while β angle (0.9° ± 16.7°) moves to lower values than the unbound (9.6° ± 16.3°) (**Figure 5B**). Both trends show that full-length H1 keeps the linker DNA collapsed in our simulations, consistent with previous experimental measurements [27]. In particular, we found that both α (20.0° ± 16.7°) and β (5.2° ± 16.3°) distributions in GH1-nucleosome simulations are in between unbound and full-length H1 results. This demonstrates that while globular H1 by itself can bring the linker DNA together, the disordered domains will significantly reinforce this converging effect.

**Figure 5:**
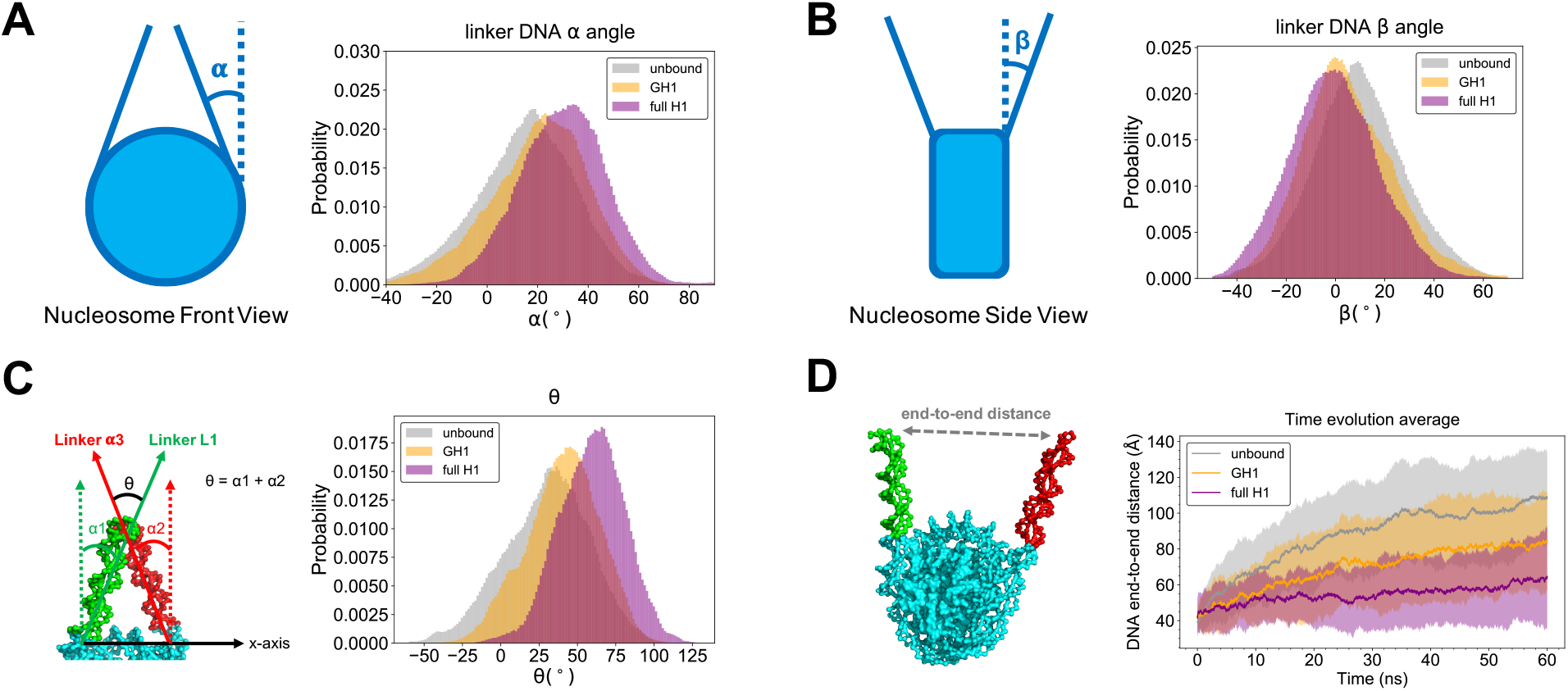
Linker DNA arms are more converged with full-length H1 than globular H1 or unbound. The definition and histogram of α, β, and θ angles are shown in **(A, B, C). (D)** The time evolution of DNA end-to-end distance, where solid lines with shaded regions represent average values with standard deviations. The same set of legends is used in all the subset figures above (gray: unbound nucleosome; yellow: GH1-nucleosome; purple: full-length H1-nucleosome).

Analyses of individual linker DNA arms allow us to measure their relative conformations and distances to describe the compaction of chromatosome with further accuracy. Here we define an angle θ = α1 + α2 between the two linker DNA arms and plot its distribution (**Figure 5C**). Its histogram demonstrates that the θ of GH1-nucleosome (40.1° ± 23.5°) only increases by a small amount compared with the unbound nucleosome result (30.2° ± 27.0°). While the binding of full-length H1 increases θ significantly (58.6° ± 21.4°). This comparison reveals the disordered domains are more crucial than the GH1 in compacting chromatosome’s structure.

We also calculated the end-to-end distance between the tips of linker DNA, and plotted its average and standard deviation over all runs along the simulation time (**Figure 5D**). We found in absence of H1, the linker DNA arms start to separate from each other after 20 ns. The end-to-end distance reaches ~ 100 Å at the end of simulations. Globular H1 slows down the linker DNA’s separation but cannot fully prevent it. The end-to-end distance still rises to ~ 80 Å by the end of simulations. On the other hand, the binding of full-length H1 remarkably inhibits this separation, as found in previous experiments [27]. The end-to-end distance does not show any apparent increasing trend beyond fluctuations. These results demonstrate that globular H1 alone is not sufficient to fully compact the chromatosome as found in previous experiments, highlighting the crucial role of disordered domains.

As a more comprehensive comparison between our simulation results and cryo-EM experiments, we also ran simulations of the H1.5ΔC50 system, another chromatosome with H1.5 linker histone, where the last 50 residues on CTD were removed, following Bednar *et al*. [27] (see Figure S7 for its sequence and structure, the simulation details are elaborated in Supporting Information). The statistics of all the DNA metrics used above (α, β, and θ angles and end-to-end distances) of H1.5ΔC50 are very similar to the full-length H1.0 results (Figure S8). This similarity of DNA dynamics reveals that H1.5ΔC50, even though belonging to a different H1 subtype and having a shorter CTD, also leads to relatively compact linker DNA geometry and dynamics. This outcome further indicates that the function of H1 disordered domains is to restrict linker DNA.

### H1 NTD/CTD are tethered to both linker DNA arms

We next computed a regional contact map for H1 NTD/CTD and DNA (**Figure 6**), using the same definitions as in **Figure 4** and Table S1. For H1 CTD, the GH1-proximal C1, C2, and C3 regions have higher contact probabilities (**Figure 6B**, 3rd-5th column), while the GH1-distal regions form very minimal contacts with DNA (**Figure 6B**, 6th-10th column). A similar trend is found for NTD, where the GH1-proximal N2 region is much more tightly bound than the distal N1 region (**Figure 6B**, 1st-2nd column). Both CTD and NTD have closely bound regions with the dyad DNA on N2, C1, and C2 (**Figure 6B**, 2nd-4th column, 2nd row). On the other hand, CTD prefers binding with L1-DNA via its C1 and C3 regions (**Figure 6B**, 3rd and 5th column, 1st row), while NTD is particularly bound with α3-DNA via the N2 region (**Figure 6B**, 2nd column, 3rd row). This contact map elucidates how the full-length H1 is steadily bound with DNA near the entry-exit site: the two disordered domains, especially the GH1-proximal parts, act as “hands” of GH1 to grasp the dyad and linker DNA. Such an entanglement between H1 disordered domains and DNA precludes GH1 from escaping far away from the dyad region and confines linker DNA dynamics.

**Figure 6:**
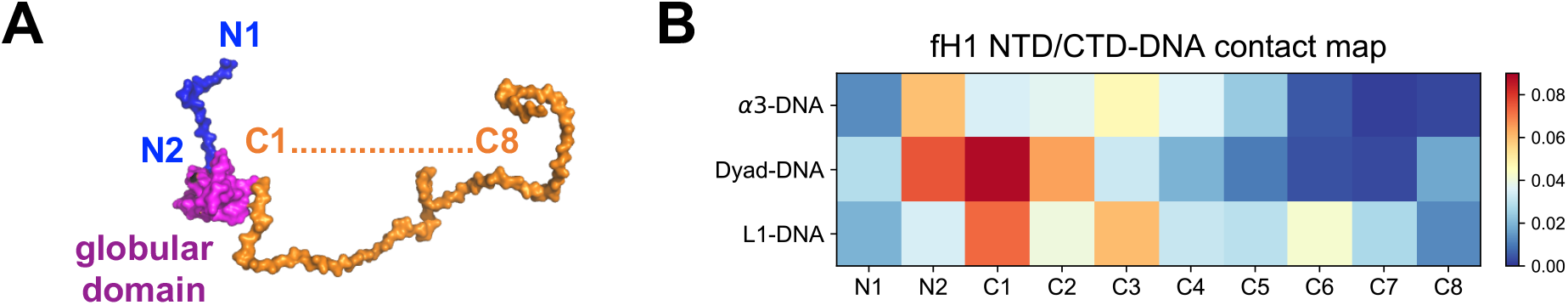
H1 NTD and CTD are tightly bound with DNA mainly via the residues proximal to the globular domain. **(A)** H1 NTD (blue) is divided into two regions (N1-N2, from GH1-distal to GH1-proximal). H1 CTD (orange) is divided into eight regions (C1-C8, from GH1-proximal to GH1-distal). **(B)** shows the contact map of DNA with NTD and CTD. The horizontal axis represents NTD (N1-N2) and CTD (C1-C8) regions. The vertical axis and color bar are the same as in **Figure 4B**.

We also observed that the contact probability of the GH1-distal region of the CTD (especially C4-C8) is much smaller, indicating that it is only transiently bound to DNA. Interestingly, the entire NTD and CTD remain highly disordered in the simulation (see Figure S9). This result agrees with previous NMR experimental results [37], which may explain absence of H1 tails in almost all chromatosome X-ray crystal structures.

## Discussion

In this work, we investigated binding dynamics of the H1-nucleosome complex using extensive computer simulations using a state-of-art protein-DNA model. Our focus was on the regulatory effect of H1 disordered domains on the chromatosome structure. By quantitative comparisons among different molecular systems, we found that H1 disordered domains, especially GH1-proximal regions, compact chromatosome structure by constraining the conformation and dynamics of linker DNA arms and GH1. By contrast, without the disordered domains, the binding affinity between the H1 globular domain and DNA is notably reduced. In this case, linker DNA arms become more separated, and GH1 may even escape away from the nucleosome, resulting in a more “open” chromatosome conformation.

Previous studies suggested that H1 can bind with a nucleosome on- or off-dyad to regulate distinct chromatin higher-order structure [25]. A recent review [22] pointed out that H1-nucleosome binding modes depend on H1 species and experimental conditions, and the resulting chromatosome structures should be viewed as an ensemble. Interestingly, our simulations not only replicate H1-nucleosome on/off-dyad binding modes found in previous experiments and all-atom simulations, but also discover new H1 binding modes far from the dyad. One commonly observed new binding mode in our simulation corresponds to GH1 completely escaping from dyad and reaching close to the acidic patch on histone core. Our findings suggest that the disordered domains play a balancing role to prevent the chromatosome from being too rigid or too flexible. The resulting structural plasticity of the chromatosome enables H1’s rapid interconversion among different modes, including binding/unbinding to the nucleosome, which, in turn, may allow timely remodeling and adjustment of chromatin compaction levels.

In the “zigzagging ladder” model of chromatin folding, H1 acts as the “rung” between nucleosome particles. Based on our findings, we propose that H1 NTD/CTD may affect the location and strength of the connection between H1 and neighboring nucleosomes, thus regulating nucleosome array organization and chromatin higher-order structure. Compared with the globular domain, H1 disordered domains are more prone to various PTMs [20]. These PTMs would enable many important biological functions, including not only chromatin folding [51], but also apoptosis [52] and DNA transcription [53]. In particular, the phosphorylations on H1 CTD have been proved to change the secondary structure of CTD and regulate chromatin condensation level [54,55]. We expect that the phosphorylation will drive the dynamics of H1 to a moderate level between the two systems in this study (globular H1 and full-length H1) because its binding affinity to the linker DNA would be weaker than the former, but stronger than the latter.

We also observed that GH1 can even escape from the dyad region and move towards the acidic patch (**Figure 3**). The reason this behavior is both novel but also potentially biologically meaningful, is that it might explain the mechanistic basis for decades-old observations that *in vitro*, H1 can reposition nucleosomes without the use of ATP or remodelers [56,57]. The spacing function of H1s remains mysterious, but our observation of H1 breaking symmetry at the center of the nucleosome might subtly lower the energetic barrier at the pseudo-dyad, allowing nucleosomal looping in advance of sliding and repositioning along the DNA fiber. Our contact analyses in **Figure 4** reveal that α2 and α3 helices are the major GH1-dyad binding regions, no matter whether H1 disordered domains are present or not. Thus, it will be promising for further experiments to mutate some key residues in these regions, such as S49, K53, K69, and K74, to test how they affect this H1 escaping-dyad motion.

Shortly after the observation of nucleosome ladders by Hewish and Burgoyne [58], coupled with classic EM experiments visualizing regularly spaced beads on a string by Woodcock [59] and the Olinses [60], early workers in the chromatin field [61] discovered the still inexplicable phenomenon of species-specific nucleosome repeat lengths. There NRLs were thought to derive from species-specific H1 variants. Our work here provides a testable theoretical framework to explore how species-specific H1 residue changes over evolutionary time [62], in the NTD, the CTD and the GD might subtly alter the motions of H1 we observed in this study, thereby potentially contributing to the global spacing of nucleosomes. We note with excitement that the advent of high-speed AFM [63–67] provides precisely the kinetic handle needed to complement H1 fast dynamics observed in FRAP studies [5,68] and H1 off/on dyad classic steady-state biochemistry experiments [69,70] glean insights into linker histone biology as it relates to chromatin spacing and folding. A comprehensive review of powerful experimental tools for probing linker histones in chromatin environment can be found in Melters *et al*. [71], where the authors successfully visualized the dynamics of nucleosome arrays with H1.5 in real time by high-speed AFM.

One limitation of our study is the lack of individual H1 NTD/CTD’s functions. Previous FRAP experiments found deletion of the longer and less conserved CTD reduces H1-nucleosome binding affinity to a much greater extent than removal of NTD [39]. Hence, it would be interesting to computationally model an H1-nucleosome system with only NTD or CTD and investigate their independent effects in regulating chromatosome structure and dynamics. Meanwhile, for different variants, H1 NTD and CTD have much less conserved sequences and distinct lengths. Our study already shows that H1.0 and H1.5ΔC50 have similar structural properties, but it would be more interesting to conduct a systematic investigation on how the sequence and length of H1 disordered domains regulate chromatosome structure and dynamics.

## Methods

### Hybrid coarse-grained model for the H1-nucleosome complex

To investigate binding dynamics of the H1-nucleosome more efficiently and accurately, we modeled this large molecular complex (~ 950 protein AA + 193 DNA bp) with a coarse-grained protein force field AWSEM [48] and a DNA force field 3SPN.2 [50], which is a different computational approach from previous landmark atomistic MD studies of nucleosome systems [72,73]. Inspired by the funneled energy landscape theory [74], AWSEM uses three beads (*C*_*α*_, *C*_*β*_ and *O*) to represent one residue and adopts both physical and bioinformatic potentials to account for amino acid interactions. AWSEM has been applied to study many different types of proteins [49,75–77], including histones [78,79] successfully. Similarly, 3SPN.2 uses three sites –phosphate, deoxyribose sugar and nitrogenous base –to represent a nucleotide and was carefully calibrated by structural and thermodynamic properties of single- and double-strand DNA. With comparable length scale and implicit solvent assumption, these two models have been combined to study several protein-DNA systems [80–83], including the nucleosome [84].

Here, we elaborate on the detailed Hamiltonian of this protein-DNA force field. For protein-protein interactions, we used a strategy similar to Zhang *et al*. [84], which is mostly consistent with the original AWSEM but also introduces two modifications: an explicit Debye-Hückel (DH) electrostatic interactions between charged *C*_*β*_ atoms (+1 for arginine and lysine, −1 for aspartic acid and glutamic acid), and a weak Gō potential with fine-tuned parameters for the entire histone octamer to bias it towards the crystal structure. Additionally, we applied the newly developed AWSEM-IDP force field [49] to model the disordered H1 CTD and NTD. We reduced the helical potential weights only for these disordered domains to avoid artificial propensity to form helices. We also performed extensive atomistic simulations for the structural segments of these disordered domains and used the resulting trajectories to bias local tail segments in AWSEM simulations, similar to Lin *et al*. [85] (see Supporting Information and Figure S1-S3 for details of atomistic simulation, sanity checks and related structure bias setup). DNA-DNA interactions followed the 3SPN.2 model.

For protein-DNA interactions, we first tested the ansatz from Potoyan *et al*. [80] and Zhang *et al*. [84], which consists of a non-residue-specific Lennard-Jones (LJ) potential and Debye-Hückel electrostatic interactions. We found these two forces are still too weak to keep a stable nucleosome structure within our standard simulation time, where DNA tends to unwrap from the histone octamer (Figure S4). In the nucleosome crystal structure, 14 arginine side chains from histone octamer insert deeply into the DNA minor grooves to further stabilize the nucleosome [2]. This important effect was not included in the previous AWSEM treatments of the nucleosome. To mimic this special interaction, we added additional site-specific Lennard-Jones forces between these pairs of arginine *C*_*β*_ and DNA phosphate beads. With this new potential, nucleosomal DNA stays wrapped around the histone core during an entire simulation (Figure S4). Detailed formulas and parameters of this nucleosome-specific potential are elaborated in the Supporting Information and Figure S4.

In summary, the formulae of the applied AWSEM-DNA force field are listed below, where *V* stands for the potential energy of each term:

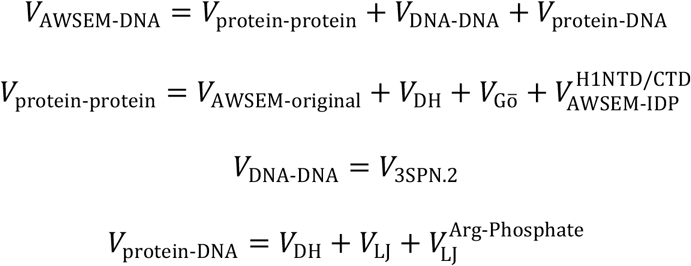

### Simulation details

All the coarse-grained simulations were performed with the open-source simulation package LAMMPS [86] (version 31Mar17), in which our modification of AWSEM-DNA was implemented [48,50,80,84]. The initial conformation of the H1-nucleosome was taken from a recent X-ray crystal structure [27] (*Xenopus Laevis* H1.0b, PDB: 5NL0). Three different molecular combinations were constructed to test the function of H1, including the nucleosome with linker DNA and (1) without H1 (unbound); (2) with the H1 globular domain; (3) with the full-length H1 (**Figure 1**). Note that the disordered H1 NTD and CTD are missing in the crystal structure. Hence we used MODELLER [87] (version 9.19) to generate their initial conformations. Tails of core histones were not included in this study, because most of them are located far away from H1 or the nucleosome dyad — the main region of interest in this study, and unlikely to interact with H1 or linker DNA. Moreover, including these disordered tails would have slowed down the convergence of our simulations.

We set the simulation time step to 5 fs and used non-periodic shrink-wrapped boundary conditions. The Langevin thermostat was applied, with the damping parameter = 500 fs. We used a parameter set to mimic electrostatic interactions in 150 mM NaCl solution, which is close to a physiological cellular environment (see Supporting Information for detailed parameters). With five different random initial velocity distributions, we heated up and annealed the system shortly (500 ps heating from 300 K to 330 K, 500 ps annealing from 330 K to 300 K) to generate five different initial configurations. Then for each initial configuration, we ran 10 independent simulations for 60 ns at 300 K and constant volume, each with different initial velocity distributions. In effect, for each molecular system we obtained simulation trajectories summing up to 3 μs, a very long timescale considering the significantly faster exploration rate in our coarse-grained simulations compared with atomistic ones.

## Analysis

### Linker DNA geometry

We used several different metrics to quantify the conformation and dynamics of linker DNA arms. We computed αand βangles between the linker DNA arms and the vertical dyad axis to quantify compaction of the chromatosome particle (see **Figure 5A-B** for graphical illustrations), following similar definitions from Bednar *et al*. [27] and Woods *et al*. [88] Vectors representing linker DNA are defined as center of mass (COM) of the beginning bp pointing to COM of the terminal bp. The vector representing the vertical dyad axis is defined as COM of bp at the center of nucleosome bottom pointing to the dyad. The “front” and “side” plane for angle αand βare defined as parallel and perpendicular to the nucleosome core disk. Positive αindicates linker DNA arms bend “inward” while positive βmeans bending “outward”. Similarly, we also defined and calculated angle θ and end-to-end distance between the linker DNA to quantify their relative geometry (see **Figure 5C-D** for graphical illustrations). We used VMD to conduct all the analyzes mentioned above.

### Protein-DNA regional contact map

We computed regional contact maps between H1 and DNA near the entry-exit site to describe their binding sites. We first divided H1 and DNA into small regions based on their secondary structure elements or location (see Table S1 for detailed region definitions). A pair of protein-DNA beads in coarse-grained representation will be determined as “contact” if their inter-bead distance is smaller than a predetermined threshold (= 8 Å, around 1.5 times of protein + DNA site radius for excluded volume effects). Based on this definition, we computed the contact probability by counting contact number within a certain H1/DNA region and normalizing it with the total number of protein-DNA bead pairs:

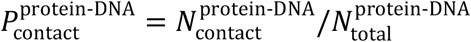

## Abbreviations

NTD/CTD: N/C terminal domain;
PTMs: post-translational modifications;
GH1: H1 globular domain;
COM: center of mass;
AWSEM: associated memory, water-mediated, structure and energy model

## Acknowledgements

Garegin Papoian’s efforts were supported by the Amazon AWS Machine Learning Research Award. Yamini Dalal’s work is supported by NIH. Computational resources are provided by the Deepthought2 HPC cluster at the University of Maryland.

