## Supporting Information for "Binding Dynamics of Disordered Linker Histone H1 with a Nucleosomal Particle"

### H1 NTD/CTD Atomistic Simulations

Particularly for the full length H1-nucleosome system, we performed atomistic simulations of the disordered NTD and CTD, and used them as bioinformatic local structural bias in the subsequent coarse-grained (CG) simulations. To simulate the relatively large CTD (99 residues) efficiently and obtain local structural information, we separated its sequence into six short overlapping segments and ran individual atomistic simulations for each segment, as done in Lin *et al.*[1] The 24-residue NTD is much shorter so we simulate its entire structure. See Figure S1 for their sequences and detailed segmentation.

We used replica-exchange molecular dynamics to enhance sampling when simulating these two intrinsically disordered regions. As for the force field, we used the recently developed a99SB-disp[2] for protein and the modified TIP4P-D[3] for water. These two force fields are specially optimized for simulating IDPs and prove more accurate than many other atomistic force fields. The temperatures of all replicas were determined by the T-REMD server.[4] The temperature range is 300 - 400 K and the target exchange probability is 0.25, generating 30 - 60 replicas at different temperature for each simulated system. All the atomistic simulations were performed with GROMACS[5] (version 2018.2) package, with time step of 2 fs, periodic boundary conditions and particle mesh Ewald method for long-range electrostatics. The initial structures (same as the CG simulation) were solvated in a dodecahedronal box. Then sodium and chloride ions were added into the solvent to neutralize the system and create a 150 mM physiological salt concentration.

The systems first underwent a short energy minimization of 100 ps using the steepest descent algorithm at 300 K. The NVT equilibration for 500 ps and NPT equilibration for 1 ns were performed subsequently at the unique temperatures of each replica, both with position constraints and Berendsen temperature or pressure coupling methods. Then we performed the production runs (100 ns for NTD and 30 ns for CTD segments), adding up to 1.8 - 3 $\mu$s total simulation time of all the replicas for each system. The trajectories at 300 K were used for following analyses and structural bias for CG simulations, excluding the first 20 ns of NTD simulation and 5 ns of CTD simulations for equilibration. The resulting NTD/CTD structures have very low ordered secondary structure propensity (Figure S2) and extended global size (Figure S3). These disordered features agree with previous studies in general, proving the sanity of using these structures to bias subsequent CG simulations.

### AWSEM-IDP Potential for H1 Disordered Domains

As mentioned in the main text, we used AWSEM-IDP[6] to model H1 disordered domains, with some recent modifications. Here we elaborate the details.

#### Fragment Memory

We used the atomistic simulation trajectories mentioned above to bias the local structure of H1 NTD and CTD in CG simulations. This structural bias is called "fragment memory" and its formula is:

$$V_{\mathrm{FM}} = -\lambda \sum_{m} \omega_m \sum_{ij} \gamma_{ij} \exp[-\frac{(r_{ij} - r_{ij}^{m})^2}{2\sigma^2}] \tag{S1}$$

where $\omega_m$ represents the weight for each memory $m$. Detailed definitions of other parameters and significance of this potential can be found in previous AWSEM literature.[6,7]

The atomistic simulation trajectories were clustered by the simple linkage algorithm in GROMACS with RMSD = 2.5 Å as threshold. The representative structures of all the clusters are then used as fragment memory. The weights for each memory are calculated based on the logarithm of cluster size based on the Boltzmann equation, and normalized to be on the same scale with weights ($\omega = 1$) of other globular proteins with single structure as memory in the system, *i.e.* for memory from $\mathrm{cluster}_m$, its weight $\omega_m$ is:

$$\omega_m = \log(\mathrm{size}_m) / \sum_{j}^{N_{\mathrm{cluster}}} \log(\mathrm{size}_j) \tag{S2}$$

#### Sequence-Specific Helical Propensity

$V_{helical}$ is a term in responsible for alpha helices formation in AWSEM. In the original version of AWSEM-IDP, we reduced its weight to 1.2 for the entire IDP chain. The disordered H1 NTD/CTD, however, are connected with the globular domain in the same protein chain. To solve this inconsistency, we implemented a modification for $V_{helical}$ so that AWSEM user can assign weights for each amino acid in a chain. Here, we reduced $V_{helical}$ weight to 1.0 for H1 NTD and CTD, and kept using 1.5 for all the other ordered proteins, including H1 globular domain.

#### $R_g$ Potential

We did not use the newly introduced $R_g$ potential from AWSEM-IDP in this study. Because there is no experimentally or computationally measured $R_g$ available for the H1 disordered domains, especially when bound with nucleosomal DNA.

### Electrostatic Interactions

All the AWSEM-DNA simulations use two types of electrostatic interactions: DNA-DNA and protein-DNA. There is no protein-protein electrostatics, because the contact term in the original AWSEM for protein has already taken short-range electrostatic interactions into consideration.

The Debye-Hückel style DNA-DNA electrostatics are directly from 3SPN.2 without modification:

$$V_{\mathrm{elec}}^{\mathrm{DNA-DNA}} = \sum_{n_{\mathrm{elec}}}^{i<j} \frac{q_i q_j e^{-r_{ij}/\lambda_D}}{4\pi\epsilon_o \epsilon(T,C) r_{ij}} \quad r_{ij} < r_c \tag{S3}$$

where $q_i$ and $q_j$ are the charge on site $i$ and $j$, $r_{ij}$ the distance between these two sites, $\lambda_D$ is the Debye length, $\epsilon_o$ is the dielectric permittivity of vaccum, $\epsilon(T,C)$ is the dielectric permittivity of solution. $\lambda_D$ and $\epsilon(T, C)$ are both dependent on ionic concentration and temperature. In this study, we set ionic concentration = 150 mM, temperature = 300 K and $r_c = 50$ Å to mimic DNA-DNA electrostatics in the physiological cellular environment. The detailed formulae of $\lambda_D$ and $\epsilon(T, C)$ and values of other parameters in DNA-DNA interactions can be found in Hinckley *et al.*[8]

Similarly, the protein-DNA electrostatics are also modeled in Debye-Hückel style in a non-sequence-specific manner:

$$V_{\text{elec}}^{\text{protein-DNA}} = \sum_{n_{\text{elec}}}^{i<j} \frac{q_i q_j e^{-r_{ij}/\lambda_D}}{4\pi\epsilon_o \epsilon r_{ij}} \quad r_{ij} < r_c \tag{S4}$$

where all the parameters have the same definition as in Eq. S3. Here we set $\lambda_D = 10$ Å, $\epsilon = 78$, $r_c = 40$ Å, which results in proper electrostatics in a 150 mM NaCl solution.

### Nucleosome Specific Arginine-Phosphate Potential

To prevent nucleosomal DNA unwrapping from the histone octamer core, we applied an additional Lennard-Jones (LJ) potential between certain $C_\beta$ beads from arginine residues on histones and phosphate beads from DNA (both in CG representation). Note that this force is only applied to 14 protein-DNA pairs to mimic the effect that some arginines are deeply inserted into DNA minor grooves.[9] See Figure S4A for the residues and base pairs involved in this force. Its formula is the standard 12/6 LJ potential:

$$V_{\text{LJ}}^{\text{Arg-Phosphate}} = \sum_{\text{pair}}^{n_{\text{pairs}}} 4\epsilon[(\frac{\sigma}{r_{\text{pair}}})^{12} - (\frac{\sigma}{r_{\text{pair}}})^{6}] \quad r < r_c \tag{S5}$$

where $r_{pair}$ is the $C_\beta$-phosphate inter-bead distance for a certain pair of arginine-DNA, $\epsilon$ is strength of the potential, $\sigma$ is finite distance at which the potential is zero, and $r_c$ is the cutoff distance. After fine tuning the parameters, we eventually set $\epsilon = 5$ kcal/mol, $\sigma = 5$ Å and $r_c = 15.5$ Å. With this parameter set, this nucleosome-specific LJ potential provides stronger protein-DNA attraction (see Figure S4B for energy comparison with other protein-DNA interaction terms) so that nucleosomal DNA can keep wrapping around the histone core. This potential is implemented as a new fix style in LAMMPS as "fix/lj/cut", inspired by the pair style "lj/cut".

To test the effect of this new potential, we run some short simulations (5 ns) for the canonical nucleosome without H1 (PDB: 1KX5) before and after applying the arginine-phosphate potential. All the other setup and parameters are the same as the H1-nucleosome simulations in the main text. We found without this arginine-phosphate potential, the DNA are highly dynamic and start to unwrap from the histone core at the end of simulations in two out of five runs (Figure S4C). By contrast, with the $V_{\mathrm{LJ}}^{\mathrm{Arg-Phosphate}}$, the nucleosomal DNA keep wrapping around in all the five simulation runs (Figure S4D).

#### Representative Snapshots from 3D Spherical Coordinates

We established a set spherical coordinates ($r$, $\theta$, $\phi$) of H1 globular domain's center of mass, as shown in the main text, to quantify their dynamics. We identified all the major basins from their 2D ($\phi$, $r$) histogram and selected the representative snapshots as follows. We first estimated $\phi$ and $r$ values at the center of each basin. Then we computed the most probable $\theta$ value with similar $\phi$ and $r$, and searched for corresponding snapshot with this ($r$, $\theta$, $\phi$) as representative ones.

#### H1.5△C50 Simulations and Analyses

To further comprehensively compare our computational results with the cryo-EM data,[10] we also performed simulations for H1.5, another subtype of linker histone, binding with nucleosome. The C-terminal 50 amino acids of H1.5 in our study was deleted to be consistent with the molecule in cryo-EM experiment. Therefore this system is called "H1.5ΔC50" (see Figure S7 for its sequence aligned with H1.0 and structure).

Similar to H1.0, H1.5 also has disordered NTD and CTD. Therefore, we first ran atomistic simulations for H1.5ΔC50 NTD/CTD structural segments. Then we simulated the H1.5ΔC50-nucleosome complex by AWSEM-DNA force field, with previous atomistic simulations as structural bias for disordered domains. All the simulation details are the same as in H1.0 case, except that we only performed 8 independent runs (total 480 ns in CG time scale), limited by computational resource and speed. Then we performed the same linker DNA analyses and compared with the cryo-EM experiments (Figure S8). We found our results are in general quantitatively consistent with Bednar *et al.*

Table S1: **H1 region definitions for contact analysis**

| Domain | Regions | $AA/BP_{start}$ | $AA/BP_{end}$ | Length (AA/BP) |
|---|---|---|---|---|
| H1NTD | N1 | 1 | 12 | 12 |
| | N2 | 13 | 24 | 12 |
| GH1 | $\alpha$1 | 28 | 38 | 11 |
| | L1/$\beta$1 | 39 | 46 | 7 |
| | $\alpha$2 | 47 | 57 | 11 |
| | L2 | 58 | 62 | 5 |
| | $\alpha$3 | 63 | 78 | 16 |
| | $\beta$2 | 81 | 87 | 7 |
| | $\beta$3 | 90 | 95 | 6 |
| H1CTD | C1 | 98 | 109 | 12 |
| | C2 | 110 | 121 | 12 |
| | C3 | 122 | 133 | 12 |
| | C4 | 134 | 145 | 12 |
| | C5 | 146 | 157 | 12 |
| | C6 | 158 | 169 | 12 |
| | C7 | 170 | 181 | 12 |
| | C8 | 182 | 196 | 15 |
| DNA | L1 | 1 | 23 | 23 |
| | dyad | 87 | 107 | 21 |
| | $\alpha$3 | 171 | 193 | 23 |

Table S2: $\phi$-$r$ **basin definitions and population percentages**

| Basin ID | $r_{min}$ (Å) | $r_{max}$ (Å) | $\phi_{min}$ (°) | $\phi_{max}$ (°) | Population % |
|---|---|---|---|---|---|
| g1 | 60 | 75 | 0 | 150 | 33.1 |
| g2 | 30 | 45 | 60 | 140 | 8.5 |
| g3 | 55 | 75 | -100 | -140 | 7.9 |
| g4 | 40 | 60 | -80 | -20 | 8.0 |
| g5 | 100 | 120 | -60 | -30 | 0.9 |
| f1 | 60 | 75 | 0 | 150 | 53.3 |
| f2 | 30 | 45 | 60 | 140 | 11.7 |
| f3 | 55 | 75 | -100 | -50 | 4.0 |

```
      H1.0 N-terminal Domain:
      .         .         .
      1         11        21
      MAENSAATPAAKPKRSKALKKSTD
      ~~~~~~~~~~~~~~~~~~~~~~~~
      1                      24

      H1.0 C-terminal Domain:
      .         .         .         .         .         .         .         .         .         .
      98        108       118       128       138       148       158       168       178       188
      ADEGKKPAKKPKKEIKKAVSPKKVAKPKKAAKSPAKAKKPKVAEKKVKKVAKKKPAPSPKKAKKTKTVKAKPVRASKVKKAKPSKPKAKASPKKSGRKK
Seg0  ~~~~~~~~~~~~~~~~~~~~~~~~
      98                     121
Seg1                 ~~~~~~~~~~~~~~~~~~~~~~~~~
                     113                     137
Seg2                                 ~~~~~~~~~~~~~~~~~~~~~~~~
                                     129                    152
Seg3                                                ~~~~~~~~~~~~~~~~~~~~~~~~
                                                    144                    167
Seg4                                                               ~~~~~~~~~~~~~~~~~~~~~~~
                                                                   159                   181
Seg5                                                                             ~~~~~~~~~~~~~~~~~~~~~~~~
                                                                                 173                    196
```

Figure S1: **H1 NTD (residue 1-24) and CTD (residue 98-196) sequences used in the simulations.** Residues with positively and negatively charged side chains are labeled in green and red respectively. The SPKK repeating motifs in CTD are underlined. Segments for atomistic simulations are represented by ∼.

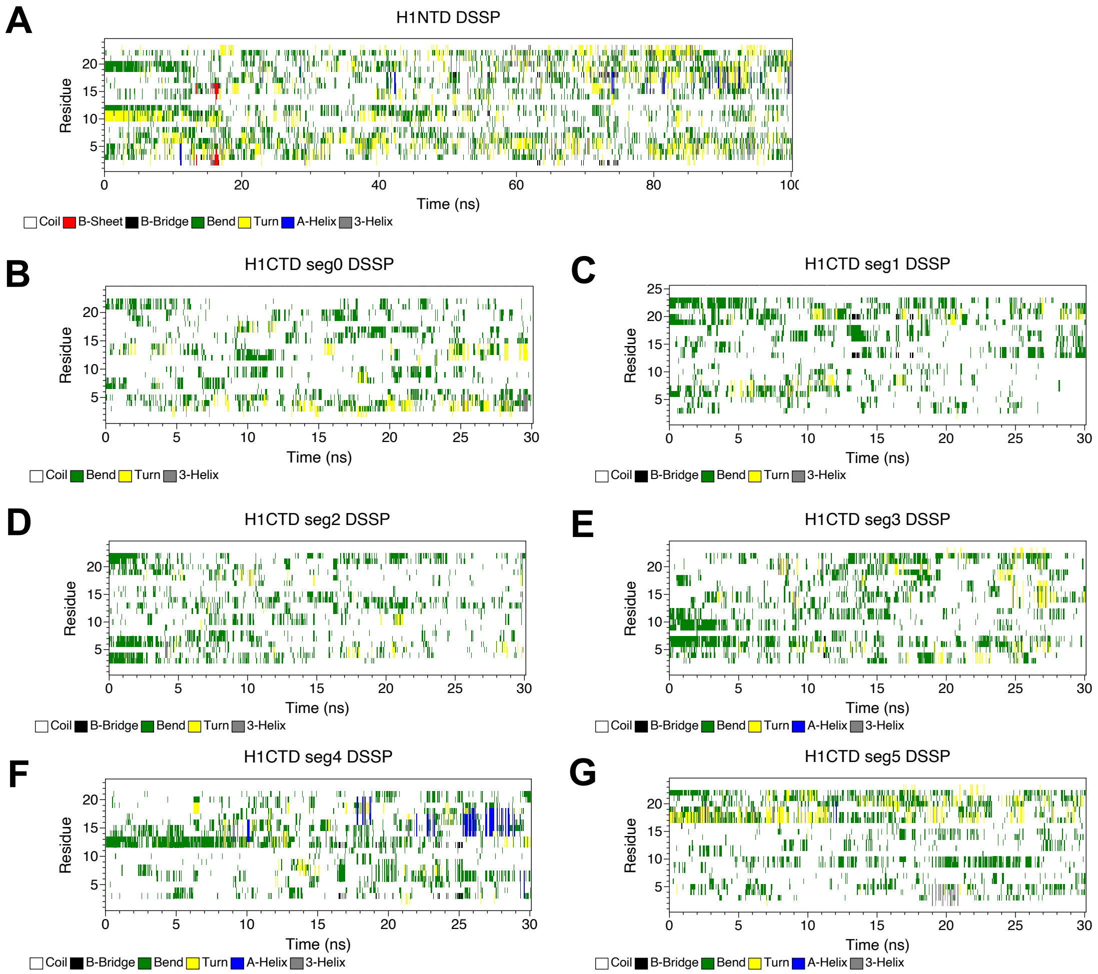


Figure S2: **H1 NTD/CTD have disordered secondary structure in atomistic simulations.** Secondary structure elements for each residue (vertical axis) along the simulations (horizontal axis) are plotted for H1 NTD (A) and CTD (B-G) segments. Color code represents different types of secondary structure. Analyses were performed using DSSP[11] in GROMACS.

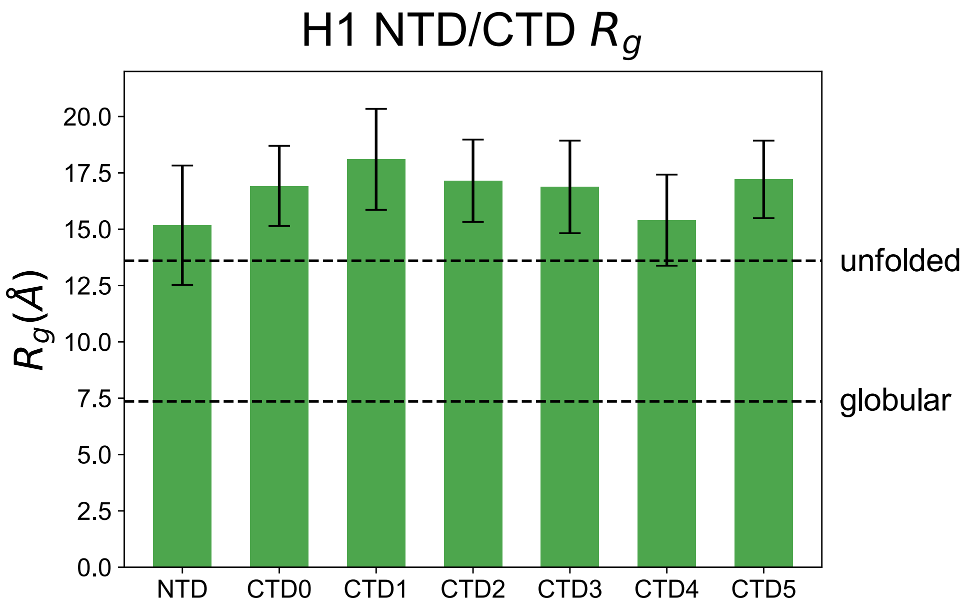


Figure S3: **H1 NTD/CTD are extended in atomistic simulations.** Radius of gyration ($R_g$) were computed for all the H1 NTD/CTD segments. The average and standard deviation of $R_g$ are shown as bar plot with error bars. The theoretical $R_g$ for globular protein[12] and unfolded random coil[13] with the same number of residues ($\sim 24$) are plotted as horizontal dashed lines.

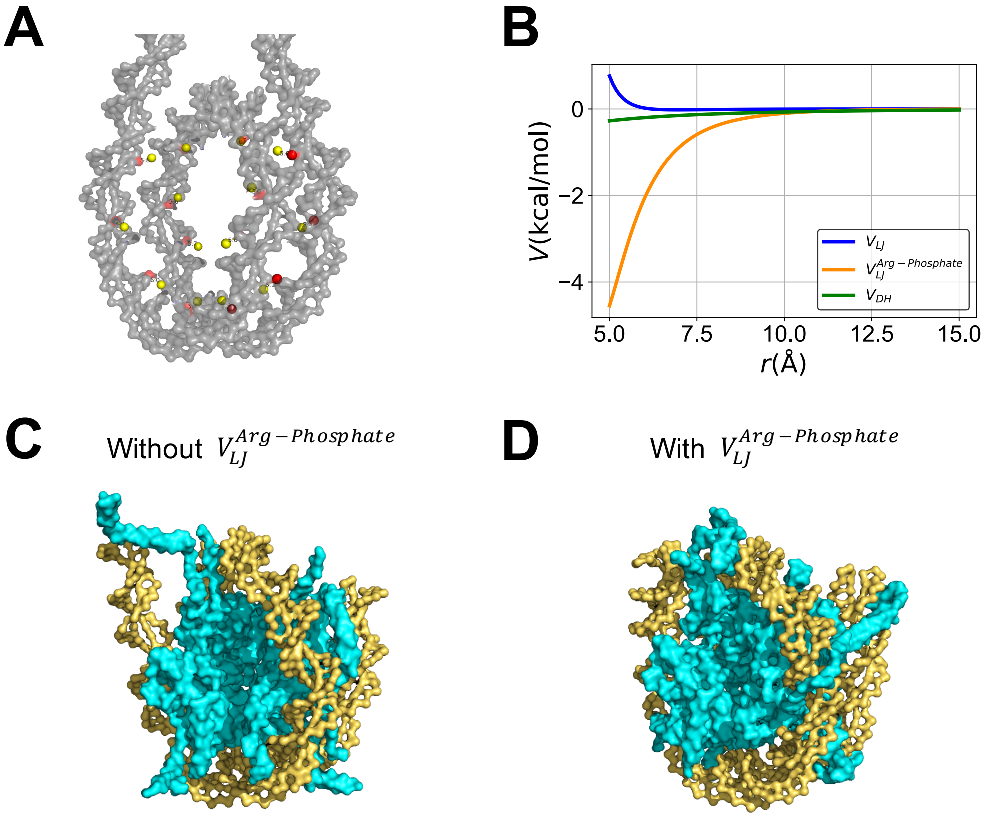


Figure S4: **A special Lennard-Jones potential is applied between protein-DNA to avoid nucleosomal DNA unwrapping.** (A) shows the locations of all 14 pairs of arginine $C_\beta$ (yellow) and DNA phosphate beads (red). Nucleosomal DNA are shown in gray, while the histone core and H1 are not shown for clarity. (B) Different protein-DNA energy terms $V$ as a function of inter-bead distance $r$ with the parameter set used in this study. The new potential ($V_{\mathrm{LJ}}^{\mathrm{Arg-Phosphate}}$) creates a significant energy barrier ($\sim$ 3 - 4 kcal/mol) from $r = 5$ Å to $r = 10$ Å, stronger than the standard LJ ($V_{\mathrm{LJ}}$) and Debye-Hückel ($V_{\mathrm{DH}}$) terms. The representative final snapshots of the test canonical nucleosome simulations without (C) and with $V_{\mathrm{LJ}}^{\mathrm{Arg-Phosphate}}$ (D) illustrate the effect of this new potential to avoid DNA unwrapping (gold: DNA; cyan: histone).

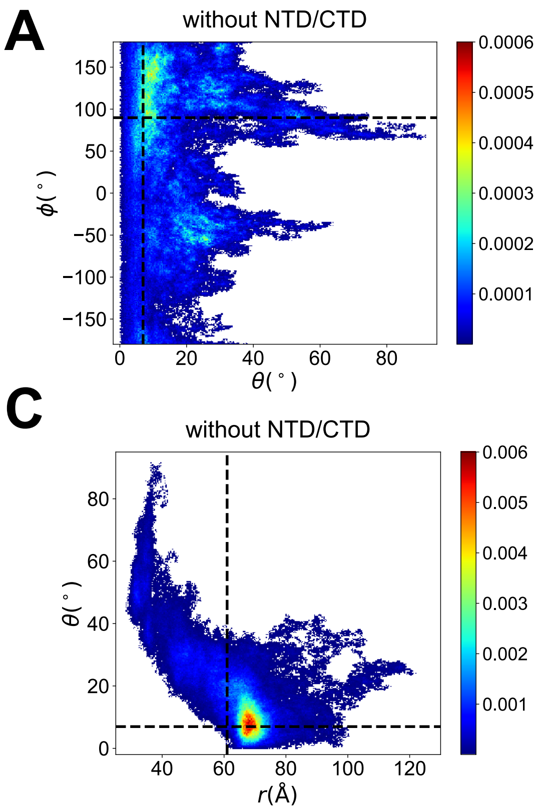


Figure S5: **Additional 2D histograms of the 3D coordinates for GH1 COM** (A-B): ($\phi$, $\theta$) without (A) and with disordered domains (B). (C-D): ($\theta$, $r$) without (C) and with disordered domains (D). The corresponding values of the chromatosome crystal structure[10] (with GH1 bound on dyad) are labeled with dashed lines.

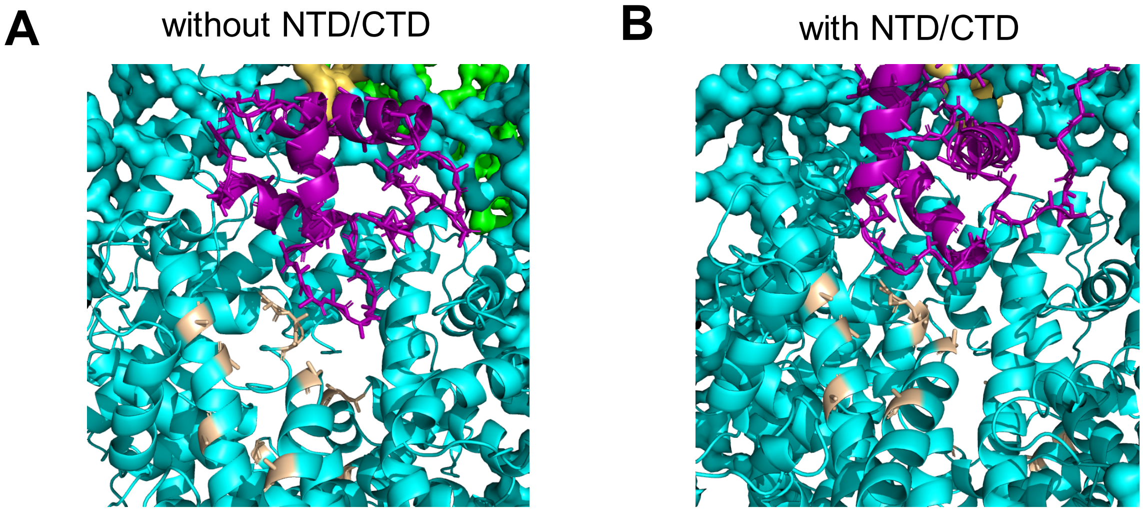


Figure S6: **Zoomed-in snapshots in which the GH1 is proximal to the histone core acidic patch**. GH1 (purple) moves close to the acidic patch (brown) on the histone core in both systems without (A) or with (B) the H1 disordered domains. These two figures as example are taken from the g2 and f2 representative snapshot in the main text. All the "sticks" representation only show a small part of the amino acid side chains because of the coarse-grained nature of our AWSEM-DNA model.

**A**

```
H1.0              1 MAENS---AATPA------AKPKRSK------ALKKSTDHPKYSDMILAA     35
                    |:|.:   .||||      ||.|.:|      |.|:....|..|::|..|
H1.5DC50          1 MSETAPAETATPAPVEKSPAKKKATKKAAGAGAAKRKATGPPVSELITKA     50

H1.0             36 VQAEKSRSGSSRQSIQKYI-KNHYKVGENADSQIKLSIKRLVTSGALKQT     84
                    |.|.|.|:|.|..:::|.: ...|.|.:| :|:|||.:|.||:.|.|.||
H1.5DC50         51 VAASKERNGLSLAALKKALAAGGYDVEKN-NSRIKLGLKSLVSKGTLVQT     99

H1.0             85 KGVGASGSFRLAK---------------ADEGKKPAKKPKKEIKKAVSPK    119
                    ||.||||||:|.|               |.:.||||....|:.|||...|
H1.5DC50        100 KGTGASGSFKLNKKAASGEAKPKAKKAGAAKAKKPAGATPKKAKKAAGAK    149

H1.0            120 KVAKPKKAAKSPAKAKKPKVAEKKVKKVAKKKPAPSPKKAKKTKTVKAKP    169
                    |..|     |:|.|||||..|  .||||||     ||||
H1.5DC50        150 KAVK-----KTPKKAKKPAAA--GVKKVAK-----SPKK-----------    176

H1.0            170 VRASKVKKAKPSKPKAKASPKKSGRKK    196

H1.5DC50        177 ---------------------------    176

N-Terminal Domain; Globular Domain; C-Terminal Domain
```

**B**

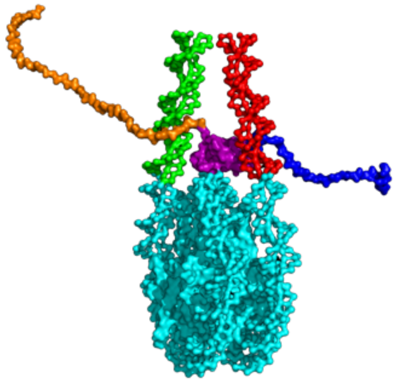

Figure S7: **H1.5ΔC50 sequence and structure**. (A) Sequence alignment of H1.0 (196 AA) and H1.5ΔC50 (176 AA) analyzed with EMBOSS NEEDLE web server.[14] The color code indicates different domains. The identical, similar and different amino acids are labeled by "|", ".", and ":". (B) H1.5ΔC50-nucleosome complex structure. The color code is the same as in the main text.

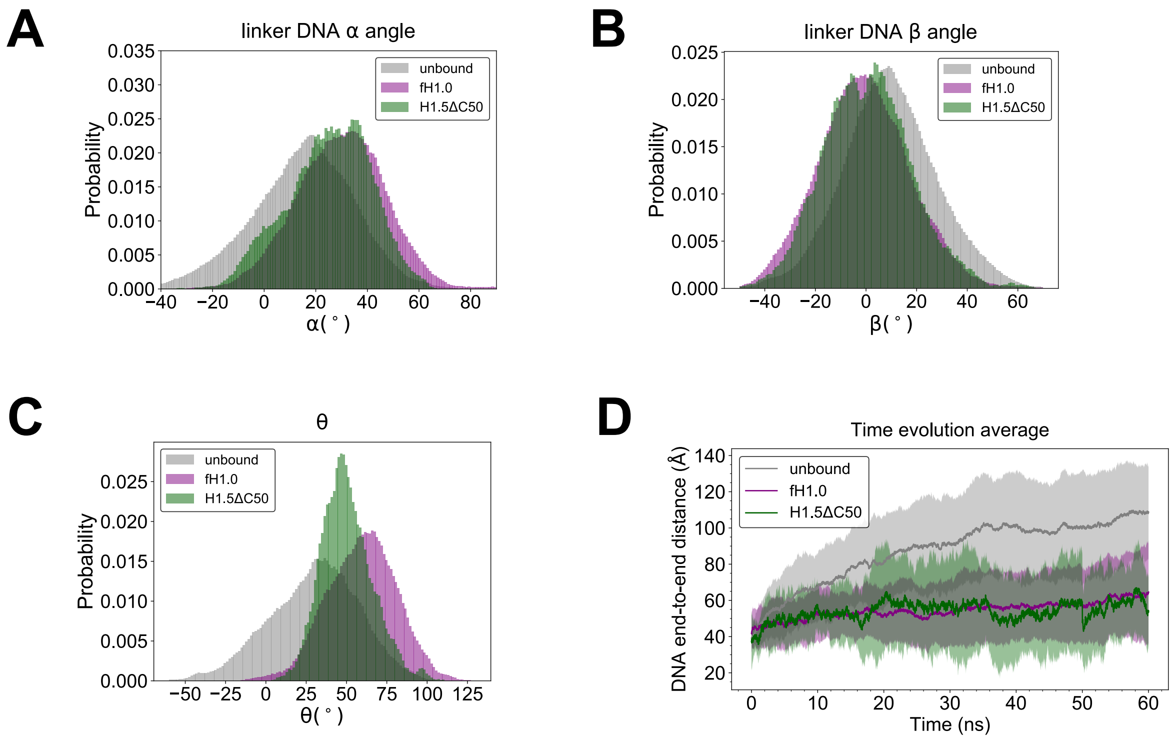


Figure S8: **H1.5ΔC50 DNA conformation and dynamics are consistent with the previous cryo-EM study.** The DNA related metrics $\alpha$, $\beta$, $\theta$ and end-to-end distance for unbound nucleosome, full-length H1.0-nucleosome and H1.5ΔC50-nucleosome are plotted using the same definition as in the main text. (A-B) qualitatively agree with the same measurements in Figure 1E of Bednar *et al.*[10]

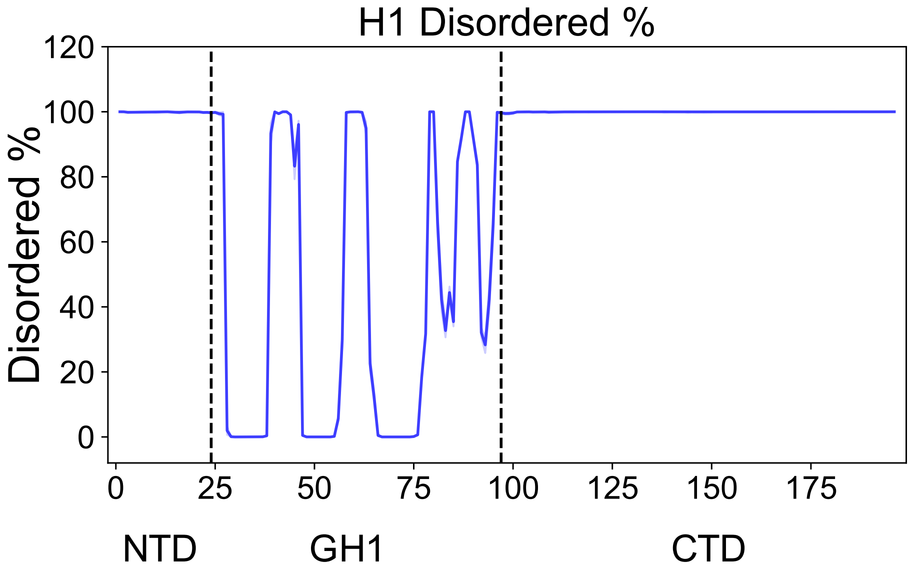


Figure S9: **H1 remains disordered in nucleosomal context.** The disordered probability at each residue is defined as the percentage that "turn" or "coil" occurs. Secondary structure elements in this analysis were determined by STRIDE[15] in VMD.
